# Acylation of MLKL impacts its function in necroptosis

**DOI:** 10.1101/2023.08.19.553906

**Authors:** Apoorva J. Pradhan, Shweta Chitkara, Ricardo X. Ramirez, Viviana Monje-Galvan, Yasemin Sancak, G. Ekin Atilla-Gokcumen

## Abstract

Mixed lineage kinase domain-like (MLKL) is a key signaling protein of necroptosis. Upon activation by phosphorylation, MLKL translocates to the plasma membrane and induces membrane permeabilization which contributes to the necroptosis-associated inflammation. Membrane binding of MLKL is initially initiated by the electrostatic interactions between the protein and membrane phospholipids. We previously showed that MLKL and its phosphorylated form (pMLKL) are *S*-acylated during necroptosis. Here, we characterize acylation sites of MLKL and identify multiple cysteines that can undergo acylation with an interesting promiscuity at play. Our results show that MLKL and pMLKL undergo acylation at a single cysteine, C184, C269 and C286 are the possible acylation sites. Using all atom molecular dynamic simulations, we identify differences that the acylation of MLKL causes at the protein and membrane level. Through systematic investigations of the *S*-palmitoyltransferases that might acylate MLKL in necroptosis, we showed that zDHHC21 activity has the strongest effect on pMLKL acylation, inactivation of which profoundly reduced the pMLKL levels in cells and improved membrane integrity. These results suggest that blocking the acylation of pMLKL destabilizes the protein at the membrane interface and causes its degradation, ameliorating necroptotic activity. At a broader level, our findings shed light on the effect of *S*-acylation on MLKL functioning in necroptosis and MLKL-membrane interactions mediated by its acylation.

## INTRODUCTION

Different types of programmed cell deaths follow complex and interconnected signaling networks and play different roles in cellular homeostasis and pathological conditions.^1, 2^ Necroptosis is one such type of programmed cell death that is characterized by cell swelling, disruption of plasma membrane and release of intracellular contents resulting in extensive inflammatory activity.^3, 4^ Although these morphological characteristics are similar to those in necrosis, which is not regulated, necroptosis has a distinct molecular mechanism. The inflammatory nature of necroptosis is due to membrane rupture and leakage of intracellular contents and is triggered by the activation of the tumor necrosis factor (TNF) receptors via TNF family of cytokines.^5^ Binding of TNF alpha (TNFα) to TNF receptor recruits multiple adaptor proteins, one of which is receptor interacting protein kinase 1 (RIPK1). This event can signal a cell survival pathway or a cell death pathway such as apoptosis when caspase-8 activity is high. Sustained presence of RIPK1 and decreased caspase-8 activity initiates necroptosis where RIPK1 forms a complex with RIPK3, followed by autophosphorylation of RIPK3.^6^ This complex results in the phosphorylation of mixed lineage kinase domain-like (MLKL) which then oligomerizes and translocates to the plasma membrane causing membrane rupture. These key interactions that result in the eventual membrane permeabilization make MLKL the executioner protein of necroptosis.^7–10^

MLKL consists of an N-terminal four-helix bundle domain (4HB) with two additional helices (brace domain) connecting the 4HB to the C-terminal pseudokinase domain.^7^ The 4HB consists of multiple positively charged amino acids that can interact with the negatively charged phosphatidylinositol phosphates (PIPs) at the plasma membrane.^11, 12^ This protein-lipid interaction is essential for MLKL binding to the plasma membrane. The interactions between the C-terminal helix of the brace with the rest of the helices can inhibit the functioning of MLKL in necroptosis.^13^ The helices of the brace are responsible for oligomerization of MLKL and also maintain contacts between the 4HB and the pseudokinase domain after phosphorylation, communicating the conformational changes within the pseudokinase domain to the 4HB.^12–14^ Work by Wang *et al.* and Dondelinger *et al.* have suggested that the membrane disruptive activity of MLKL results from the interactions between the positively charged residues in the 4HB and the negatively charged PIPs species present in the membrane leaflet by liposomal models.^8, 11^ Similarly, Su *et al.*, using reconstituted membranes, have suggested that the 4HB can directly insert into the membranes, causing membrane disruption.^13^

The C-terminal of MLKL is a pseudokinase.^7^ Multiple studies implicate the pseudokinase domain functioning as a ‘switch’ to regulate necroptotic signaling, downstream of MLKL phosphorylation with several charged residues within the domain essential for driving the oligomerization and translocation to the plasma membrane.^15, 16^ A study elucidated the sequence of MLKL activation by proposing a model of initial dimerization of the pseudokinase domain after phosphorylation by RIP3 followed by oligomerization via the brace region.^17^ Overall, the functioning of MLKL as the executioner of necroptosis involves multiple features and dynamic changes that warrants further investigations.

Previous studies from our lab revealed additional protein-membrane interactions that mediate MLKL binding to the plasma membrane during necroptosis. We showed that MLKL and pMLKL are *S*-acylated by very long chain fatty acids (fatty acids with ≥ 20 carbons^18^) during necroptosis and that this acylation increases their membrane binding and exacerbates membrane permeabilization in this process^19^. However, the biochemical characterization of MLKL acylation and, importantly, the implications of acylation on MLKL functioning in necroptosis remained unknown. In this work, we first systematically studied the cysteines that can be possible targets of acylation in necroptosis. Using all atom molecular dynamics simulations, SwissPalm predictions and cysteines with previously unaccounted functions, we prioritized cysteines that are likely to interact with the lipid bilayer since palmitoyl transferases are membrane-associated. Mutagenesis experiments followed by acyl-PEG exchange (APE) suggested that C184, C269 and C286 can undergo acylation in necroptosis. We then comprehensively characterized the involvement of plasma membrane-localized palmitoyl transferases in the acylation of MLKL and pMLKL during necroptosis. We demonstrate that zDHHC21 activity has the most profound effect on MLKL and pMLKL acylation. Inactivating zDHHC21 disrupted the acylation of pMLKL, ameliorated plasma membrane and, intriguingly, decreased the overall levels of pMLKL, most likely through increased protein degradation. Altogether, these results provide new insights into the interactions that regulate the role pMLKL in necroptosis.

## RESULTS

### C184, C269 and C286 of MLKL are acylated in necroptosis

Our previous efforts to understand protein acylation in necroptosis showed that MLKL and pMLKL undergo a mono *S*-linked acylation during necroptosis downstream of plasma membrane translocation.^19^ The human MLKL has 10 cysteines across the 4HB, the brace and the pseudokinase domain, with C18, C24, C28 and C86 within the 4HB; C184 in the brace helix and C269, C286, C415, C437 and C448 within the pseudokinase domain (**Figure 1A**).^20^ The four cysteines in the 4HB, C18, C24, C28 and C86 are involved in oligomerization of MLKL and induction of necroptosis.^10^ Hence, they are likely buried in the oligomerization interface and are not accessible to palmitoyltransferases located at the plasma membrane. Further, studies report that C86 is the target residue of necrosulfonamide which inhibits MLKL oligomerization^21, 22^, supporting that C86 is not likely to interact with the plasma membrane and be available for acylation. Based on these, we decided to interrogate the remaining cysteines as acylation targets in necroptosis.

**Figure 1:**
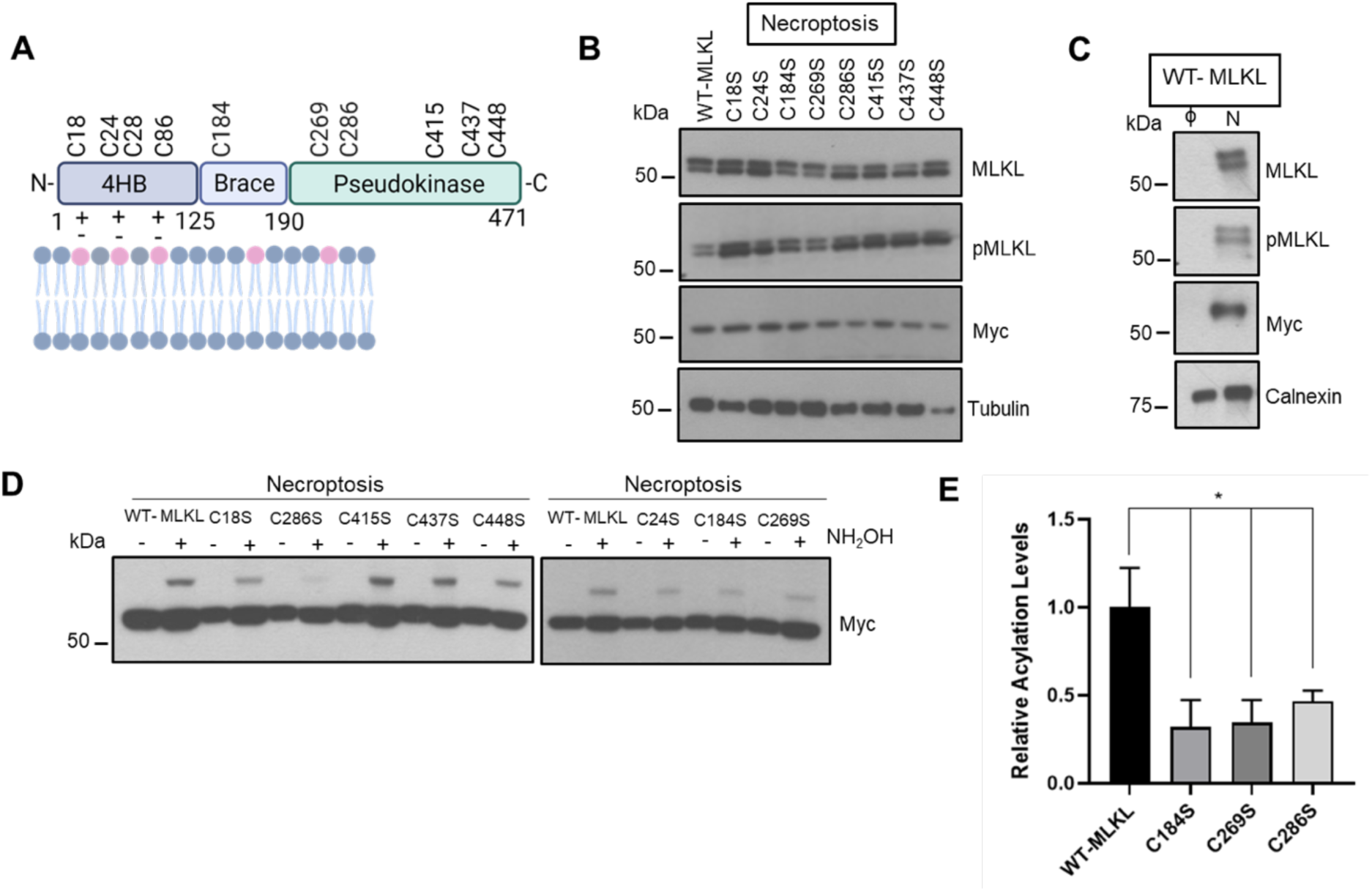
Acylation of cysteines in MLKL. **(A)** Representation of human MLKL showing the N-terminal 4-helix bundle (4HB) connected to the C-terminal pseudokinase domain via the two helix brace. 4HB spans from amino acid residues 1-125, brace helices upto residue 190 and the pseudokinase domain till residue 471.^20^ The positively charged residues in the 4HB interact with the negatively charged PIPs (shown in pink) via electrostatic interactions. The ten cysteines present in human MLKL and their location have been shown. The cysteines 184, 269 and 286 which show acylation in our experiments have been marked in red. Figure created using BioRender. **(B)** Western blot analysis of Myc-tagged WT-MLKL and the 8 constructs with C to S mutations. Necroptosis was induced in HT-29 cells following treatment with 1 μM BV6, 25 μM zVAD-fmk and 10 ng/μL TNFα and collected after 3h. Each of the constructs expresses the Myc-tagged MLKL and also undergoes necroptosis. MLKL and pMLKL show up as two bands with the Myc-tagged form being the top band and endogenous forms being the bottom bands. **(C)** WT- MLKL control and necroptotic cells were treated with C20 alkFA and fractionated. The membrane fraction was subjected to CuAAC to attach biotin-azide followed by enrichment on neutravidin resin. The samples were blotted for MLKL, pMLKL and Myc. Calnexin was used as loading control. **(D)** C184S, C269S and C286S show profound reduction in acylation as compared to WT-Myc in necroptosis. Acyl PEG exchange with N-ethylmaleimide (NEM), hydroxylamine (NH_2_OH) and methoxypolyethylene glycol maleimide (mPEG-Mal) was performed on necroptotic WT-MLKL and cysteine mutant cells. The conditions without NH_2_OH (represented by -) are controls for presence of acylation. The samples are blotted for myc. **(E)** Quantification of the extent of acylation for C184S, C269S and C286S with respect to control WT-myc. The extent of acylation is calculated by dividing the background-subtracted intensities of migrated Myc-tagged MLKL relative to non-migrated Myc-tagged MLKL for WT-Myc and C184S, C269S and C286S (using Fiji). The relative values are calculated by normalizing values for the mutants with respect to the values obtained for Myc-tagged WT-MLK. Data represent the mean ± 1 SD; n=3. * represents *p* < 0.05

We first used all-atom molecular dynamic simulations to investigate the interaction of murine MLKL with a membrane model based on the lipid content of HT-29 cells.^23^ The simulations identified different protein-bound conformations with the 4HB or the pseudokinase domain interacting with the lipid membrane; the 4HB was found to have deeper membrane insertion, in agreement with previously published reports of the 4HB being responsible for contact with the lipid membrane and subsequent insertion.^13^ The trajectories showing interaction of the pseudokinase domain with the membrane resulted in a different lipid landscape around the protein after the simulation reached equilibrium.^23^ Upon further examination of these trajectories, we observed C23, C379, C280 and C407 consistently have the highest occupancy in terms of percentage of time bound to the membrane (**Table S1**, highlighted in green). The corresponding cysteines in human MLKL were then selected for further investigations (C18_human_→C23_murine_, C286_human_ →C280 _murine_, C415 _human_ →C407 _murine_; Y387_human_→C379 _murine_ in MLKL, **Figure 1A, Figure S1A**). In parallel, we consulted with SwissPalm database^24, 25^ which predicts C18 and C24 to be palmitoylated. The remaining cysteines, C184, C269, C437 and C448, do not have any known role in MLKL functionality in necroptosis. Based on our simulation results, SwissPalm predictions and literature information available, we shortlisted C18, C24, C184, C269, C286, C415, C437 and C448 to be investigated for their role in acylation of MLKL in necroptosis.

We conducted mutagenesis studies to investigate the involvement of the above-mentioned cysteines in pMLKL and MLKL acylation during necroptosis. We first subcloned a commercially available MLKL sequence in a pLYS1 vector expressing a Myc tag and introduced point mutations to generate MLKL_C18S_, MLKL_C24S,_ MLKL_C184S,_ MLKL_C269S,_ MLKL_C415S,_ MLKL_C437S_ and MLKL_C448S_ constructs. These were then used to generate lentiviruses to transduce HT-29 cells and express Myc-tagged WT MLKL (WT-MLKL) and mutant forms of MLKL. We confirmed the expression of WT-MLKL and mutant forms and the ability of the HT-29 cells expressing Myc-tagged MLKL forms to undergo necroptosis by Western blotting (**Figure 1B**). Briefly, we induced necroptosis using BV6, a SMAC mimetic and a pan-caspase inhibitor zVAD-FMK followed by TNF-α we described previously.^19, 26, 27^ We collected cells after 3h of necroptosis induction and samples were analyzed using Western blotting. The two bands in Figure 1B correspond to the endogenous MLKL (lower band, MW: ∼55 kDa) and the Myc-tagged MLKL (upper band, MW: ∼57 kDa). Both WT and mutant forms of MLKL can undergo acylation in necroptosis, supporting that the mutation of Cys to Ser does not impact necroptotic signaling or MLKL functionality (**Figure 1B**). We further confirmed the ability of Myc-tagged WT-MLKL to undergo acylation during necroptosis by using a clickable C20 FA probe (**Figure 1C**). These results established that Myc-tagged MLKL can undergo acylation, similar to the endogenous MLKL in necroptosis.

We then used APE to study the role of these cysteines on the extent of acylation of MLKL. APE involves removing the acyl chains on modified cysteines with hydroxylamine to generate free sulfhydryl groups and then tagging these sulfhydryl groups with a high molecular weight polyethylene glycol (PEG) which introduces a mass shift that can be observed using SDS-PAGE^28^. This allows to calculate the extent of acylation for the mutants by comparing the levels of acylated Myc-tagged MLKL to the level of non-acylated Myc-tagged MLKL expressed in the cells, for control WT-MLKL and each of the mutants. We observed that three mutant forms of MLKL, MLKL_C184S_, MLKL_C269S_ and MLKL_C286S_, showed a significant decrease (p < 0.05, t-test) in acylation as compared to WT-MLKL (**Figure 1D-E**). Specifically, C184S and C269S mutations resulted in a 68% and 65% reduction whereas C286S mutation showed a 53% reduction in acylation of MLKL-Myc. We note that the C24S mutant also showed ∼20-30% reduction in acylation but this reduction was not statistically significant (p > 0.05).

While predominantly a single cysteine residue of MLKL is acylated during necroptosis, our mutagenesis results point to a preference of three cysteines, C184, C269 and C286, as acylation sites. C184 is situated in the brace helix of MLKL, with the other two cysteines in the pseudokinase domain^20^ with the brace in mouse MLKL found to interact with membrane lipids.^29^ This observation points to a mechanism where acylation of either of the three cysteines during necroptosis, locks the protein in a specific conformation at the membrane surface upon insertion of the acylated tail, with the other two cysteines being unmodified.

### Acylated MLKL interacts with membrane lipids

We have previously developed an all atom simulations pipeline of a complex membrane model composed of a mixture of 5 lipid species [dioleoyl-phosphatidylcholine (DOPC), dioleoyl-phosphatidylethanolamine (DOPE), cholesterol, palmitoyl-oleoyl-phosphatidylinositol 4-phosphate (PIP), palmitoyl-oleoyl-phosphatidylinositol (2,5)-bisphosphate (PIP_2_)] with the murine MLKL protein (PDBID: 4BTF).^23^ We showed that MLKL interacts with membrane lipids and leaves a specific fingerprint on the membrane local composition upon binding. Here, we used this pipeline to investigate MLKL-membrane interactions in the acylated form of the protein. Our simulations comprised of an *S*-acylated MLKL with C24 FA (a representative very long chain fatty acid) at C286 (residue 280 in murine MLKL), based on the mutagenesis results presented in Figure 1, and the same membrane model. We simulated a total of three replicas referred to as 280r1, 280r2, and 280r3 (**Figure 2**). We characterized protein-lipid interactions in acylated vs non-acylated MLKL to assess the effect of acylation in MLKL-membrane interaction.

**Figure 2:**
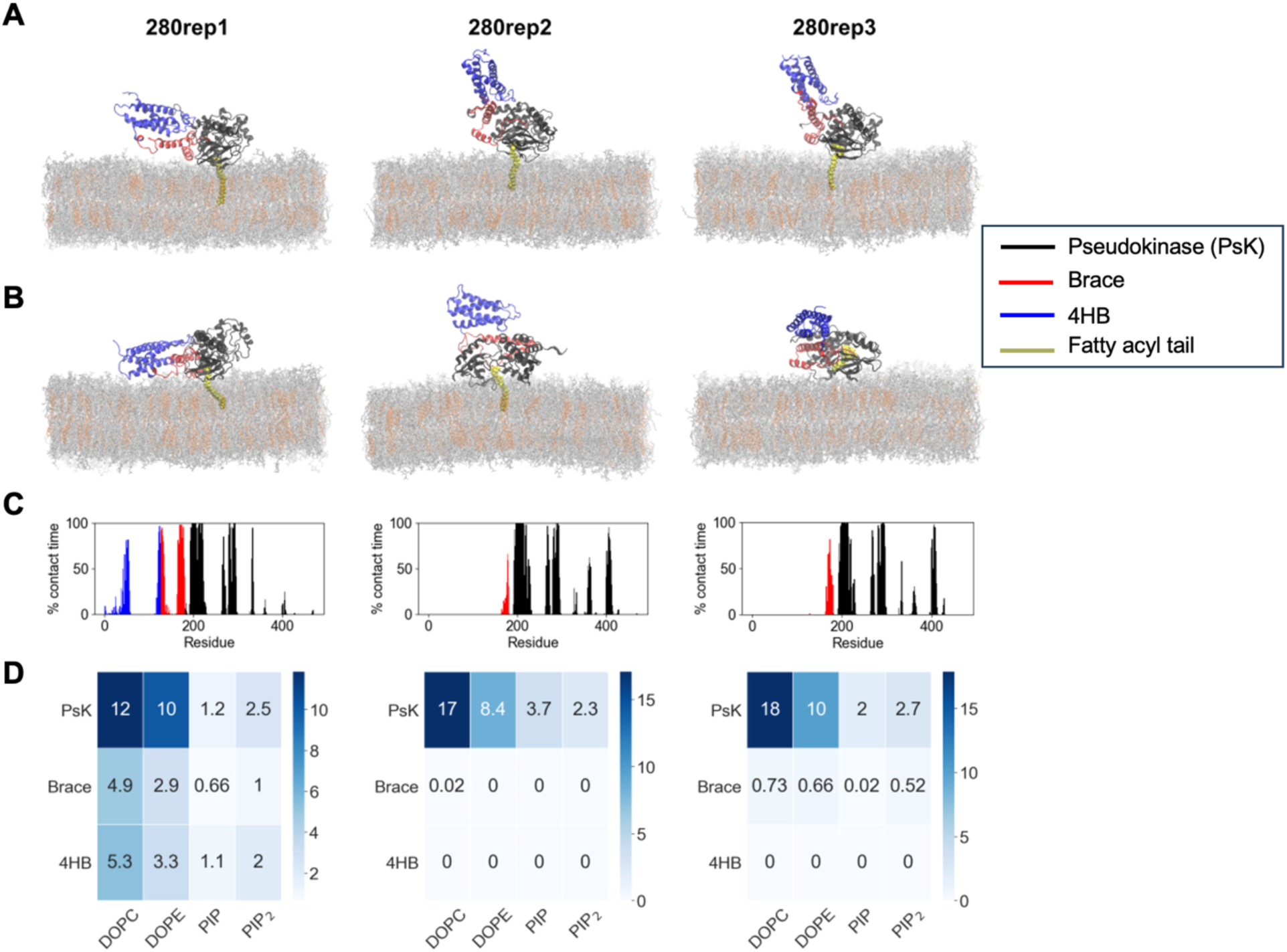
Molecular dynamics shows interaction of acylated MLKL with membrane lipids. (**A**) Position of the protein in the membrane after 50ns of equilibration, initial interaction; (**B**) after 1050ns of simulation; (**C**) Frequency of contact of each residue in the protein during the simulation; (**D**) Mean lipid distribution of the lipids within 12 Å of the protein for the last 200ns of simulation.

Figure 2A-B shows the protein starting from a bound conformation with the acylated tail partially or fully inserted into the membrane, and the final bound conformations, respectively. The lipid tail remains inserted during the full trajectory for 280r1 and 280r2, whereas it reverts to a hydrophobic pocket in the protein around 250ns in replica 280r3. The most striking difference between the acylated system (Figure 2) and that of the endogenous MLKL^23^ is the conformational change of the protein in 280r2 and 280r3. In these replicas in the acylated MLKL, 4HB (shown in blue) shifts its original position and aligns with the brace in a parallel fashion (shown in red), which did not occur in the non-acylated version. In all our simulation trajectories, we observe characteristic profiles for the protein-lipid frequency of contact depending on the specific protein domain that interacts with the membrane (Figure 2C). The conformational changes observed in the 4HB and brace conformational changes appear to result from the insertion and interaction of the acyl tail and other membrane lipids (Figure 2D). When inserted into the membrane, the 24- carbon acyl tail behaves like other lipids in the membrane and influences the lateral distribution of lipids nearby, which in turn modulates protein conformational changes on the surface. The interplay between protein and lipids is highly dynamic and can result in protein conformational changes that pull the acyl tail from the membrane temporarily, as in 280r3 (Figure 2B). It is plausible that a local lipid environment rich in VLCFAs would provide a more hydrophobic environment to stabilize the inserted acyl tail and promote membrane disruption. Alternatively, when more MLKL units are present, conformational changes observed in the simulation of the monomer may not occur simply due to steric effects.

To examine the influence of protein bound conformation and acylation on the internal structure of the membrane, we investigated the splay angle of each lipid species (**Figure S2**). We report the average angle between the fatty acid tails of lipids underneath the protein site, which provides insights on the local spatial changes of membrane lipids upon protein binding. **Figure S2A** shows a general trend of lower splay angle when the protein is acylated (280r1-280r3) compared to higher angles for the membranes with a non-acylated protein (Rep1-Rep4), reported previously^23^. Furthermore, **Figure S2B-E** shows the splay angles of PIP and PIP_2_ lipids in the binding leaflet (blue bars) are also smaller than the angles between the fatty acid tails in the non-binding leaflet (orange bars) when the acylated protein interacts with the membrane. The opposite trend is observed, in general, for the splay angles of lipids under the non-acylated protein. This indicates a preference for a closed conformation for the fatty acid tails of individual lipids right underneath the acylated protein, especially when a VLCFA is present, yet an opposite response in the lipids in the non-binding leaflet. It is plausible that this local rearrangement of lipid tails conformation increases as more protein units oligomerize at the membrane interface during necroptosis, resulting in lipid platforms with markedly different physical properties than the surrounding membrane. Such changes in the lateral packing of lipid tails will also affect membrane permeabilization.

### zDHHC21 is the major regulator of MLKL acylation

We have previously shown that the acylation of MLKL occurs at the plasma membrane downstream to its phosphorylation and oligomerization and that zDHHC5 plays a role in the acylation of MLKL in necroptosis.^19^ Specifically, inactivating zDHHC5 reduced the acylation of MLKL and pMLKL but did not completely block it, suggesting that there might be other acyltransferases at the plasma membrane that are involved in the acylation of MLKL and pMLKL during necroptosis.

The zDHHC family of palmitoyltransferases consists of 23 members residing mainly at the endoplasmic reticulum and Golgi with a few localized to the plasma membrane and the mitochondria.^30^ These enzymes can exhibit a certain degree of substrate specificity although some redundancies have also been observed. Based on the promiscuity and multiple domains impacting substrate recognition, it is difficult to predict protein substrate specificity of zDHHCs using bioinformatic tools. Here, we took a systematic approach to fully characterize the zDHHC family of proteins involved in acylation of MLKL and pMLKL in necroptosis. We focused on zDHHC members that localize at the plasma membrane, including zDHHC5, zDHHC20 and zDHHC21.^31^ We confirmed that these zDHHCs are indeed expressed in HT-29, our model cell line of necroptosis.^26^ We then knocked down zDHHC5 (as a condition known to impact acylation^19^), zDHHC20 and zDHHC21 using shRNA (Figure 3A, shzDHHC5, shzDHHC20, shzDHHC21; shRFP was used a negative control), with 70-90% efficiency (Figure 3A). We then investigated the effect of reduced zDHHC activities on the acylation of MLKL and pMLKL in necroptosis using APE. We induced necroptosis in shzDHHC5, shzDHHC20, shzDHHC21 and shRFP cells. We compared the intensities of non-acylated-MLKL and -pMLKL (lower band) and their acylated form (migrated, upper bands, indicated by black arrows, Figure 3B). We quantified the difference in acylation between the knockdowns by calculating the ratios of the migrated band intensities of MLKL and pMLKL with the respective non-migrated band intensities. As expected, we observed a significant decrease in acylation of MLKL and pMLKL in shzDHHC5 (30% and 18%, respectively). Interestingly, the inactivation of zDHHC21 had a stronger effect on acylation (60% and 55% reduction for MLKL and pMLKL, p < 0.05) during necroptosis (see Figure 3B for a representative Western blot image; quantifications of Western blots are shown in Figure 3C-D). The knockdown of zDHHC20 had no effect on acylation of MLKL and pMLKL (**Figure S3A-B,** p > 0.05, t-test) in necroptosis albeit efficient knockdown levels (Figure 3A). Consistent with the extend of reductions in acylation, inactivating zDHHC21 resulted in a strong decrease in PI uptake (∼60%, p < 0.01, compared to 40% in shzDHHC5 cells, p < 0.05) while the knockdown of zDHHC20 had no appreciable effect on membrane permeability in necroptotic cells as compared to control counterparts (Figure 3E). Overall, these results show that while multiple zDHHCs can acylate MLKL and pMLKL, zDHHC21 has the most profound effect their acylation during necroptosis and contribute to the membrane permeabilization in this process.

**Figure 3:**
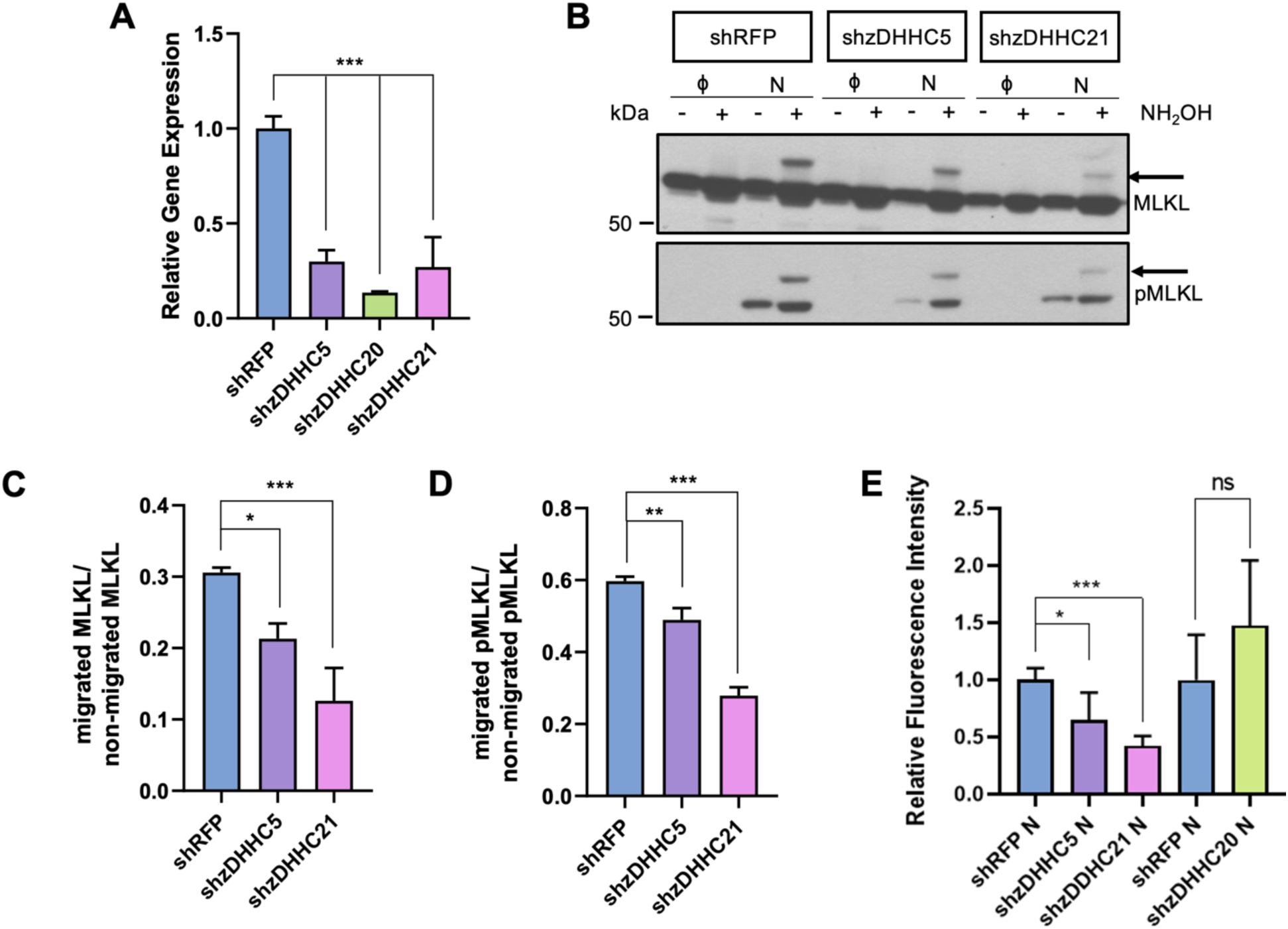
zDHHC21 has the largest effect in regulating acylation of MLKL and pMLKL in necroptosis. (**A**) Gene expression levels indicating knockdown of zDHHC5, zDHHC21 and zDHHC20. We knocked down zDHHC5, zDHHC21 and zDHHC20 using a lentiviral shRNA vector in HT-29. Relative gene expression levels are calculated as ratio of expression level of each target gene compared to the control HPRT1 expression levels. (**B**) Acyl PEG exchange with NEM, NH_2_OH and mPEG-Mal in control and necroptotic shRFP, shzDHHC5 and shzDHHC21 cells. (**C**) Quantification of acylated MLKL in shzDHHC5 and shzDHHC21 compared to shRFP. The ratio is calculated by dividing migrated band intensities of MLKL with non-migrated band intensities. (**D**) Quantification of acylated pMLKL in shzDHHC5 and shzDHHC21 compared to shRFP. The ratio is calculated by dividing migrated band intensities of pMLKL with non-migrated band intensities. (**E**) zDHHC knockdown affects membrane permeabilization. Relative fluorescence intensity represents the propidium iodide uptake of shzDHHC cells compared to control shRFP cells in necroptosis. Data represent mean ± 1 SD; n=4. * represents *p* < 0.05, *** represents *p* < 0.001.

### Reducing acylation impacts cellular levels of pMLKL during necroptosis

Our results demonstrate that reducing MLKL and pMLKL acylation via zDHHC21 inactivation restores membrane integrity during necroptosis (Figure 3E). Previous studies showed that zDHHC21 depletion ameliorated septic injury in renal reperfusion^32^ and barrier dysfunction in endothelial inflammation^33^, suggesting an important role for zDHHC21 in membrane integrity. Based on these, we decided to focus on the involvement of zDHHC21 in the fate of pMLKL, the functional form of MLKL in necroptosis. We induced necroptosis in shzDHHC21 and shRFP cells and analyzed pMLKL levels using Western blotting. To account any changes in basal expression of MLKL due to zDHHC21 knockdown, we quantified pMLKL levels relative to the total MLKL level. We observed ∼50% decrease (*p* < 0.01) in total pMLKL present in the lysate (Figure 4A) and 53% (decrease *p* < 0.001) in the membrane-localized pMLKL (Figure 4B) in shzDHHC21 cells as compared to the control shRFP cells during necroptosis. While the reduction of plasma membrane-residing pMLKL is expected upon disruption of its acylation, the overall decrease of pMLKL under these conditions is interesting and suggest that zDHHC21 activity is important for maintaining cellular levels of pMLKL during necroptosis and that reducing zDHHC21 activity decreases cellular pMLKL levels.

**Figure 4:**
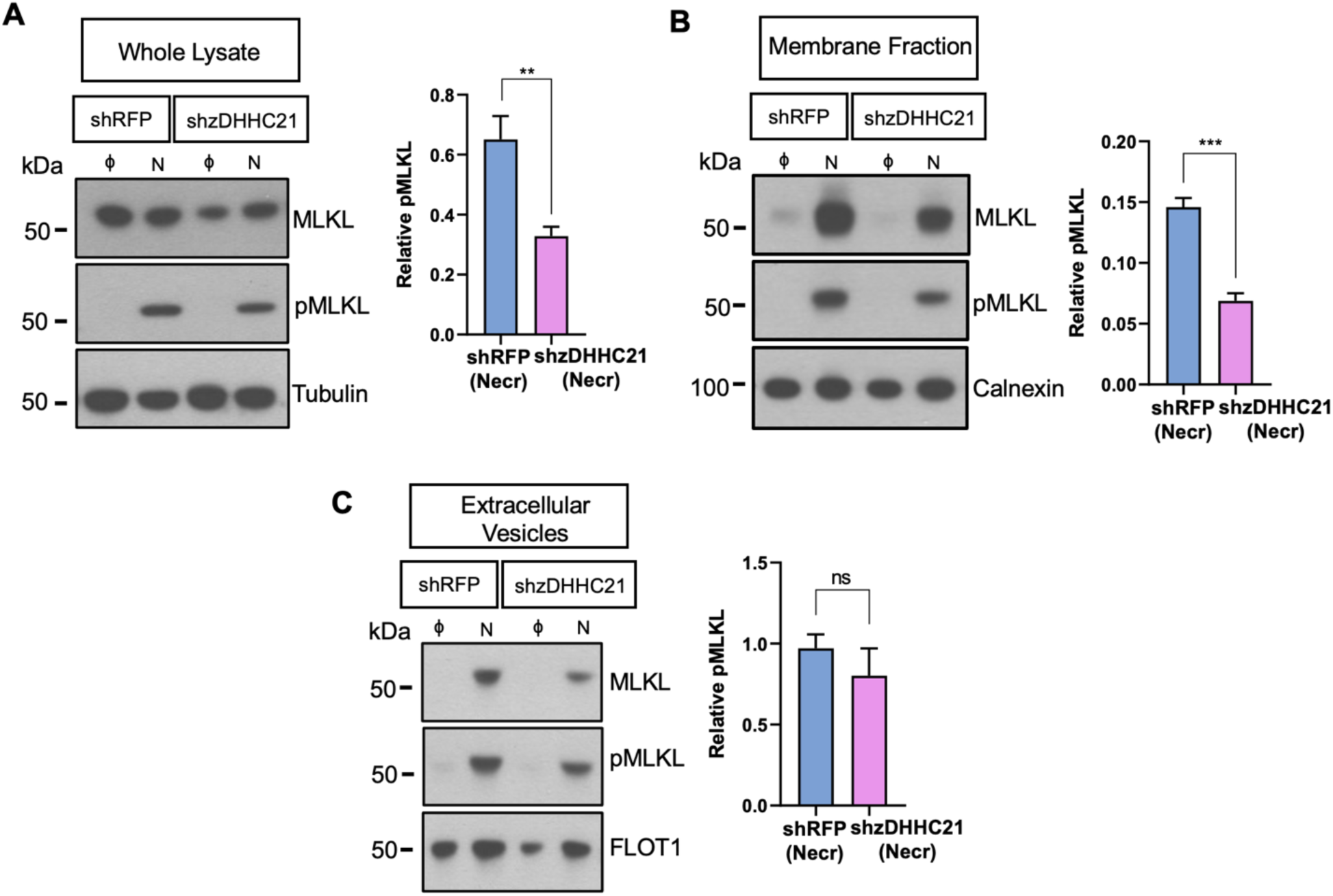
zDHHC21 regulates pMLKL levels during necroptosis. **(A)** shzDHHC21 has decreased levels of pMLKL present in the cells during necroptosis. Control and necroptotic shRFP and shzDHHC21 whole lysates were normalized based on protein amounts. The samples were blotted for MLKL and pMLKL. Tubulin was used as the loading control. The bar plot represents the quantification for the Western blot. The relative pMLKL to total MLKL levels were calculated by dividing the corrected intensities of pMLKL (normalized to tubulin) with respective normalized MLKL. Normalized MLKL for shRFP and shzDHHC21 was calculated by dividing the corrected intensities of MLKL (normalized to tubulin) with the average corrected MLKL band intensities in shRFP or shzDHHC21 control. **(B)** shzDHHC21 cells have decreased pMLKL present at the membrane during necroptosis. Control and necroptotic shRFP and shzDHHC21 cells were fractionated and normalized based on protein amounts. The samples were blotted for MLKL and pMLKL. Calnexin was used as the loading control. The bar plot represents the quantification for the Western blot. The relative pMLKL to total MLKL levels were calculated by dividing the corrected intensities of pMLKL (normalized to calnexin) with respective normalized MLKL. Normalized MLKL for shRFP and shzDHHC21 was calculated by dividing the corrected intensities of MLKL (normalized to calnexin) with the average corrected MLKL band intensities in shRFP or shzDHHC21 control. **(C)** Levels of pMLKL released from shzDHHC21 and shRFP cells remain unchanged during necroptosis. The bar plot represents relative pMLKL levels released by necroptotic shzDHHC21 and shRFP cells. The samples were normalized based on protein amounts and samples blotted for pMLKL and FLOT1. Plot represents relative protein levels calculated by normalizing the corrected intensities for pMLKL with the corrected intensities for FLOT1. Data represent mean ± 1 SD; n=3. ** represents *p* < 0.01

Several studies suggested that MLKL and pMLKL are involved in endocytosis, cellular trafficking and exocytosis during necroptosis.^34, 35^ To exclude the idea that the reduction in pMLKL we observed when blocking its acylation is due to its secretion to the extracellular environment, we investigated the levels of pMLKL levels in the extracellular vesicles when zDHHC21 is inactivated. We collected cell culture media from necroptotic and control shzDHHC21 and shRFP cells and isolated the extracellular vesicles by ultracentrifugation. Necroptotic shzDHHC21 cells showed a 33% (p<0.001) reduction in pMLKL, released through extracellular vesicles as compared to as shRFP necroptotic cells (**Figure S4A**), excluding increased secretion in shZDHHC21 cells to reduce cellular pMLKL levels. We note that the levels of released flotillin 1, a protein commonly associated with extracellular vesicles^36^, also decreased in necroptotic shzDHHC21 cells (18%, p = 0.086, **Figure S4B**) suggesting an overall reduction in extracellular components released in shzDHHC21 cells and that relative pMLKL levels remain unchanged during necroptosis (Figure 4C). Altogether, these findings clearly suggest that zDHHC21 activity plays a role in regulating cellular pMLKL levels and maintaining the integrity of the plasma membrane during necroptosis.

### Reducing zDHHC21 activity increased pMLKL degradation

Necroptosis involves a complex regulation of MLKL levels.^35^ These include lysosomal^34^ and ubiquitylation-mediated proteasomal degradation^37^. Recent studies suggest targeting proteasomal degradation of MLKL is an effective way to limit necroptotic cells death via the reduction of MLKL.^38^ Our results showing that reduced zDHHC21 activity decreased cellular pMLKL levels lead us to suspect the involvement of degradation pathways that might impact pMLKL levels when its membrane association is reduced. To probe the link between pMLKL acylation and proteasomal degradation in necroptosis, we treated shRFP and shzDHHC21 cells with (*R*)-MG132, a small molecular inhibitor of the proteasome.^39, 40^ Following treatment with (*R*)-MG132 and induction of necroptosis as we described earlier, cells were collected and analyzed using Western blotting (Figure 5A-B). We compared the effect of (*R*)-MG132 on pMLKL levels in necroptotic shzDHHC21 and necroptotic shRFP cells to assess the contribution of proteasomal degradation in pMLKL levels when its membrane association in reduced. We observed a 43% increase (*p* < 0.05) in pMLKL levels in necroptotic shzDHHC21 cells as compared to necroptotic shRFP cells (Figure 5B). These results suggest that the proteasomal degradation of pMLKL is increased when its acylation and membrane association is reduced. To support our observations on reduced acylation promoting pMLKL degradation, we used cycloheximide (CHX) which blocks protein translation^41^, a useful tool to study protein synthesis in cells^42^. shzDHHC21 and shRFP cells were treated with CHX prior to induction of necroptosis and samples were collected at 0, 3 and 5 hours of necroptosis (Figure 5C-D). The effect of zDHHC21 on the degradation of pMLKL in necroptosis is shown by comparing the band intensities of the proteins in shRFP and shzDHHC21 cells in the presence of CHX treatment for 3 and 5 hours of necroptosis, respectively (Figure 5C). We observed similar decreases in pMLKL levels after 3 hours and 5 hours of CHX treatment in necroptosis (Figure 5D) supporting our observation that reduced acylation of pMLKL increases its degradation. We note that MLKL has a long half-life (28h in HeLa cells after CHX treatment^43^) which is likely responsible for its sustained levels in this experiment. Altogether, these results suggest a mechanism by which zDHHC21-mediated acylation contributes to the retention of pMLKL at the plasma membrane during necroptosis and reducing the acylation of pMLKL targets it for degradation (Figure 5E).

**Figure 5.**
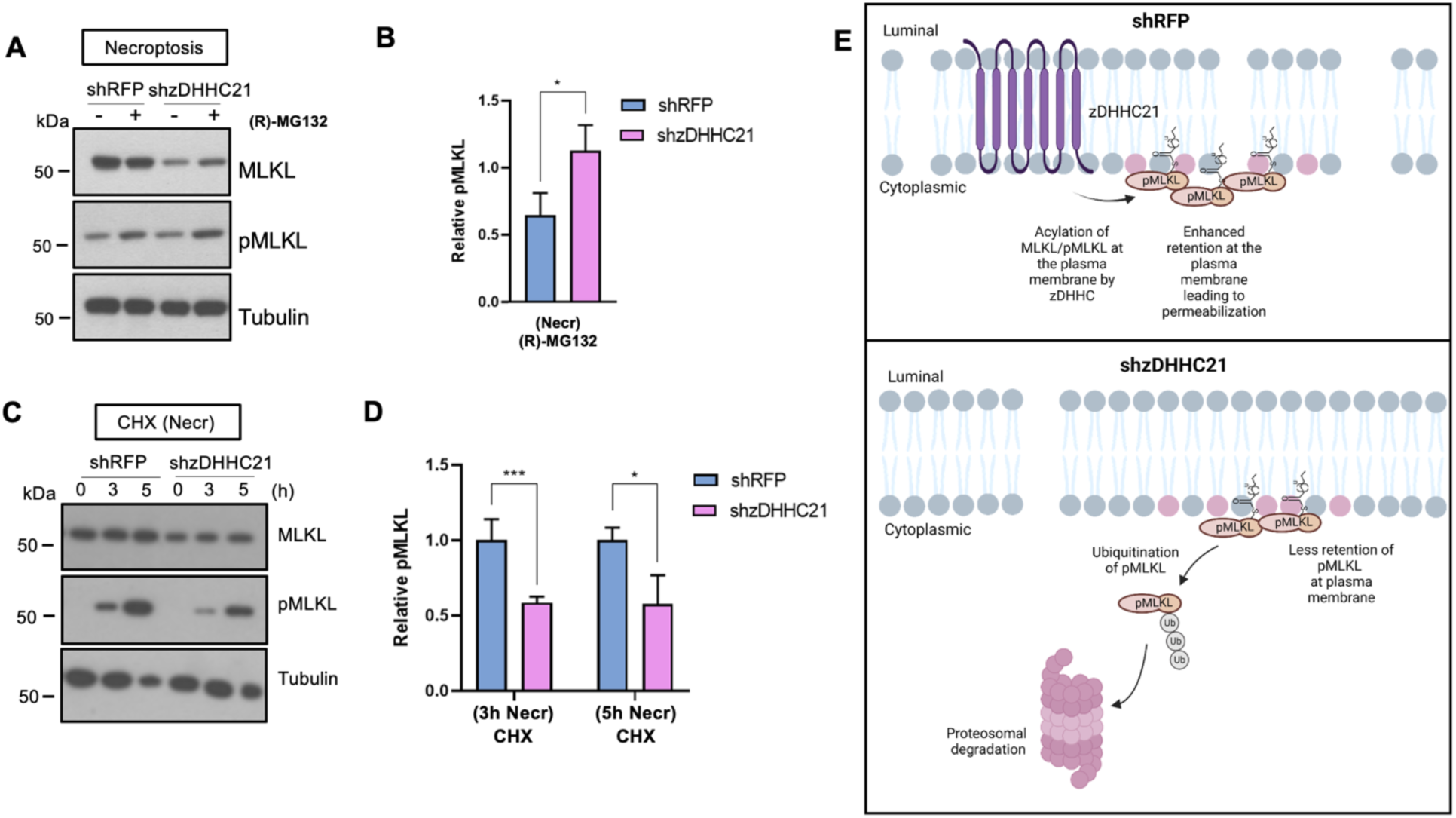
zDHHC21-mediated acylation regulates pMLKL stability during necroptosis. **(A)** Western blot of cytosolic fraction of shzDHHC21 and shRFP cells after (R)-MG132 treatment. Cells were pretreated with 10 µM (R)-MG132 for 24h, followed by 3h of necroptosis. Cells were fractionated to obtain the cytosolic proteins. Samples were blotted for MLKL and pMLKL. Tubulin was used as a loading control. **(B)** Relative pMLKL levels calculated by dividing the corrected intensities of pMLKL (normalized to tubulin) with respective normalized MLKL. Normalized MLKL for shRFP and shzDHHC21 was calculated by dividing the corrected intensities of MLKL (normalized to tubulin) with the average corrected MLKL band intensities in shRFP or shzDHHC21 control. **(C)** Western blot analysis of shRFP and shzDHHC21 in the presence of 300 μg/mL cycloheximide (CHX). Samples were collected at 0, 3 and 5 h post-necroptosis induction and prepared for Western blotting. Samples were blotted with MLKL, pMLKL and tubulin. **(D)** Quantification of pMLKL in shzDHHC21 relative to shRFP at 3 and 5 h post-necroptosis. The relative intensities are calculated by dividing the corrected intensities of each protein with the corrected intensities of tubulin and normalizing to necroptotic shRFP control. MLKL in necroptosis as a way to limit cell death before being overcome with death signal. Loss of zDHHC21 results in lesser retention of pMLKL at the plasma membranes due to reduced acylation, instead targeting pMLKL for degradation resulting in cell death rescue. Figure created using BioRender. Data represent the mean error bars represent ± 1 SD; n=3. * represents *p* < 0.05, *** represents *p* < 0.001. **(E)** A proposed model for the role of zDHHC21 during necroptosis. Cells promote degradation of pMLKL in necroptosis as a way to limit cell death before being overcome with death signal. Loss of zDHHC21 results in lesser retention of pMLKL at the plasma membranes due to reduced acylation, instead targeting pMLKL for degradation resulting in cell death rescue. Figure created using BioRender.

## DISCUSSION

Studies on MLKL has gained traction in the past few years.^44^ Other than its necroptotic function, not much is known about the physiological role of MLKL, although its involvement in endocytic trafficking has been recently reported.^35^ It has been shown that MLKL is maintained in the cytoplasm, which then clusters at the plasma membrane under necroptotic stimuli, leading to membrane disruption.^45, 46^ Previous studies have highlighted the electrostatic interactions between MLKL and the plasma membrane-associated PIP_2_ to be an important contributor to membrane binding of MLKL during necroptosis. Adding to these known interactions, we have previously demonstrated that MLKL is acylated during necroptosis. Simulation of a non-acylated MLKL with a membrane model showed PIPs are indeed recruited to the protein binding site.^23^ These results supported the dynamic interaction between MLKL and membrane lipids and suggest local membrane lipid reorganization upon MLKL binding. Here, we characterized acylation on MLKL and study the impact acylation on cellular fate of MLKL. We initially identified the acylation sites on MLKL. While the monoacylated MLKL is the predominant form, our results show that C184, C269 and C289 can undergo acylation, albeit one at a time, suggesting a promiscuity for the cysteine preferences. All atom MD simulations using an acylated form of MLKL and realistic membrane models that can mimic experimentally established interactions show a particular lipid signature upon interaction with the protein, supporting the acyl chain can cause re- structuring of the membrane lipids and contribute to membrane disruption. Comparing the trajectories of acylated MLKL and membrane to our previously reported trajectories on non-acylated MLKL^23^, we observe a conserved lipid fingerprint for the individual conformations as well as distinct local rearrangements in the presence of lipid tail in the acylated form. These interactions can result in local membrane environments with different physical properties which may affect membrane permeabilization. We believe that integrating such state-of-the-art modeling approaches with experimental data is key for generating detailed models of how acylation of MLKL contributes to membrane permeabilization at the molecular level.

We also show that while multiple S-acyltransferases can modify pMLKL, zDHHC21 has the strongest affect in acylating pMLKL during necroptosis. Reducing the activity of zDHHC21 decreases the membrane association of pMLKL and restores membrane integrity during necroptosis. Further, blocking zDHHC21-mediated pMLKL acylation promoted the degradation of pMLKL, reducing the level of this key signaling protein. Based on these results, we envision a dual role for pMLKL acylation in necroptosis. First, this acylation increases the plasma membrane binding of pMLKL and exacerbates necroptosis. Second, the acylation mediated-membrane association of pMLKL prevents its degradation and, reducing pMLKL acylation can increase its degradation and ameliorate necroptotic activity. Overall, these results establish previously unknown mechanisms that modulate pMLKL functioning during necroptosis.

## SUPPLEMENTAL INFORMATION

Supplemental Information includes 4 figures and 3 tables.

## ACKNOWLEDGMENTS

We acknowledge the support from the National Science Foundation grant (MCB1817468 to G.E.A.G.) and The Research Foundation, The State University of New York (#1181362). Simulations were run in part at the University at Buffalo’s Center for Computational Research^47^ and in the Anton2 machine. Anton2 computer time was provided by the Pittsburgh Supercomputing Center (PSC) through Grant R01GM116961 from the National Institutes of Health, specific award MCB200093P. The Anton2 machine as PSC was generously made available by D.E. Shaw Research.^48^

## AUTHOR CONTRIBUTIONS

Experiments were designed by A.J.P., V. M-G., Y.S. and G.E.A-G. The experiments were conducted by A.J.P. and S.C.Y.S. The all-atom molecular dynamic simulations were conducted by R.X.R. and V.M-G. The manuscript was written by A.J.P., R.X.R., V. M-G. and G.E.A-G. The study was directed by G.E.A-G. The authors declare no conflict of interest.

## MATERIALS

The cell lines were obtained from ATCC^®^ (HT-29 cells, HTB38, adult Caucasian female origin). For the cell culture, DMEM and penicillin/streptomycin were obtained from Corning, FBS was obtained from Sigma and trypsin (from bovine pancreas) was obtained from Millipore Sigma (Cat. # T6567-5X20UG; CAS: 9002-07-7). Plasmids were obtained from Millipore Sigma (shzDHHC5 andshzDHHC21 in pLKO.1 vectors provided as bacterial glycerol stocks, shzDHHC5: NM_015457, TRCN0000162357; shzDHHC21: NM_178566, TRCN0000144923). Plasmid pLYS1-Myc for overexpression of MLKL was obtained from Dr. Yasemin Sancak; psPAX2 was a gift from Dr. Didier Trono (Addgene plasmid, Cat. # 12260); pCMV-VSV-G was a gift from Dr. Bob Weinberg (Stewart et. al., 2003; Addgene plasmid, Cat. # 8854), shRFP (rfp_59s1c1; TRCN0000072205) was a gift by Dr. Yasemin Sancak and shzDHHC20 was a gift by Dr. Eric Witze. The QuikChange II XL Site-Directed Mutagenesis Kit was obtained from Agilent Technologies (Cat. # 200521). For plasmid extraction, the E.Z.N.A^®^ Plasmid DNA Mini Kit was obtained from Omega Bio-Tek (Cat. # D6945-01).

TNF-α (carrier free) was obtained from R&D Systems (Cat. # 210-TA/CF), BV6 was obtained from Selleck Chemicals (Cat. # S7597; CAS: 1001600-56-1) and zVAD-FMK was obtained from Enzo Life Sciences (Cat. # ALX-260-020; CAS: 220644-02-0). Propidium Iodide, Pierce^TM^ Protease Inhibitor (Cat. # A32955), Pierce^TM^ High Capacity Neutravidin Agarose (Cat. # 29202) and M- PER^TM^ Mammalian Protein Extraction Reagent (Cat. # 78501) were obtained from ThermoFisher Scientific. Puromycin (Cat. # P8833; CAS: 58-58-2), Polybrene (Cat. # TR-1003) and X-treme GENE 9 transfection reagent (Cat. # XTG9-RO) were obtained from Millipore Sigma. Cycloheximide (Cat. # 14126) was obtained from Cayman Chemicals.

Antibodies were obtained from Cell Signaling Technology (rabbit monoclonal anti-MLKL, Cat. # 14993S, RRID: AB_2721822; rabbit monoclonal anti-pMLKL, Cat. #91689, RRID: AB_2732034, rabbit monoclonal anti-Myc tag, Cat. #2278, RRID: AB_490778, rabbit monoclonal anti-calnexin, Cat. # 2679S, RRID: AB_2228381), Proteintech (rabbit polyclonal anti-Flotillin 1, Cat. # 15571-1- AP, RRID: AB_2106746), Millipore Sigma (mouse monoclonal anti*-*α tubulin, Cat. # T9026, RRID: AB_477593), Promega (goat anti-rabbit HRP conjugate, Cat. # W4011, RRID: AB_430833) and Jackson Immunoresearch Lab (goat anti-mouse HRP conjugate, Cat. # 115-035-174, RRID: AB_ 2338512).

Coomassie (Bradford) Protein Assay Kit (Cat. # 23200), Pierce^TM^ BCA protein assay kit (Cat. # 23225) and Supersignal^TM^ West Pico Chemiluminescent substrate (Cat. # 34080) were obtained from ThermoFisher Scientific. Fatty acid free BSA (Cat. # A7030, CAS: 9048-46-8), Triton^XM^ X- 100 (Cat. # T9284), sodium dodecyl sulfate (SDS) (Cat. # L3771, CAS: 151-21-3), azide-PEG3- biotin conjugate (Cat. # 762024, CAS: 875770-34-6), methoxypolyethylene glycol maleimide (mPEG) (Cat. # 63187, CAS: 99126-64-4) and N-ethylmaleimde (NEM) (Cat. # E3876, CAS: 128-53-0) were obtained from MilliporeSigma. Tris(2-carboxyethyl)phosphine (TCEP) hydrochloride (Cat. # K831, CAS: 51805-45-9) was purchased from VWR. Tris[(1-benzyl-1H-1,2,3-triazol-4- yl)methyl]amine (TBTA) (Cat. # 18816, CAS: 510758-28-8) was obtained from Cayman Chemicals.

## METHODS

### All atom molecular dynamics simulations

We conducted molecular dynamics (MD) simulations of an S-acylated version of a single MLKL protein (PDBID:4BTF) and a membrane model with 600 lipids per leaflet. The membrane was modelled after the PM with a mixture of 5 lipid species, dioleoyl-phosphatidylcholine (DOPC): cholesterol (Chol): dioleoyl- phosphatidylethanolamine (DOPE): palmitoyl-oleoyl-phosphatidylinositol-5-phosphate (POPI- 1,5): palmitoyl-oleoyl-phosphatidylinositol-(2,4)-bisphosphate (POPI-2,4) (40:32:20:4:4 mol%). Hereon after, POPI-1,5 and POPI-2,4 are referred to as PIP and PIP2. The model was built using CHARMM-GUI *Membrane Builder* ^49, 50^, and the protein was acylated with a 24 carbons lipid tail at 280C using the *Solution Builder* and in-house Python scripts. The protein was initially positioned with the 280C residue facing the membrane and the lipid tail partially inserted into the membrane. We ran three simulation replicas of the protein-membrane system with neutralizing ions for 50 ns at 310.15K using GROMACS ^51^, the CHARMM36m forcefield ^52–54^, and an integration time of 2fs. In each replica, the membrane lipids started from a different random distribution to ensure independent trajectories. The final configurations of these replicas were transferred to the Anton2 machine ^48^ to extend the simulation for 1us. The pressure was maintained at 1 bar using the Parrinello-Rahman barostat ^55^ with an isothermal compressibility of 4.5 × 10^-5^ bar^-1^, and we used the Particle Mesh Ewald methods to account for long range electrostatics ^56^.

### Cell culture and culture conditions

Human colorectal adenocarcinoma HT-29 cell (ATCC^®^ HTB-38™, adult Caucasian female origin) were cultured at 37⁰C in 5% CO_2_ atmosphere in DMEM supplemented with 10% (v/v) fetal bovine serum and 1% (v/v) penicillin/streptomycin solution. Cells were cultured for approximately 2 months and were routinely checked for mycoplasma infection. Cells were plated according to the requirement of each experiment as described below. Knockdown of zDHHC5, zDHHC20 and zDHHC21 in HT-29 cells was performed as described previously.^57^ For knockdown, cells were transduced with lentiviral particles packaged with shzDHHC5, shzDHHC20, shzDHHC21 and shRFP in pLKO.1-Puro vector and stably transduced cells were selected with puromycin. For overexpression, commercially available pLYS1-Myc vector was used and subcloned to overexpress MLKL. Lentiviral particles of overexpression system of MLKL and MLKL mutants were packaged similarly as for knockdown.

## METHOD DETAILS

### Necroptosis treatment

To induce necroptosis cell death, HT-29 cells were initially sensitized to TNF-dependent cell death pathway by SMAC mimetic BV6 (1 μM), and were co-treated with pan-caspase inhibitor zVAD-FMK (25 μM) and incubated for 30 min at 37⁰C. Cells were then treated with TNF-α (10ng/mL) and was incubated for 3 h.

### (*R*)-MG132 treatments

8×10^6^ HT-29 cells were plated in overnight. Cells were pretreated with 10 μM of (*R*)-MG132 in DMSO solution for 24h, followed by with BV6, zVAD-FMK and TNF-α treatment. Cells were collected after 3h of necroptosis and ultracentrifuged at 100,000 rcf for 45 min at 4⁰C to separate the cytosolic proteins from the membrane proteins. The cytosolic proteins were precipitated using cold 1x PBS, ice-cold methanol and ice-cold chloroform in a 1:2:1.5 ratio and samples were normalized based on protein content. Samples were prepared for Western blotting and 5 μg of sample was loaded for each protein of interest.

### Cycloheximide treatments

1.5×10^6^ HT-29 cells were plated in 6-well plates overnight. Cells were treated with 300 μg/mL cycloheximide (CHX) in DMSO solution along with BV6 and zVAD-FMK 30 minutes prior to addition of TNF-α. Cells were collected at 0, 3 and 5h following necroptosis inducted, lysed and normalized based on protein content. Samples were prepared for Western blotting and 5 μg of sample was loaded for each protein of interest.

### Propidium iodide (PI) uptake

Cells (40×10^3^) were seeded in 96-well plates overnight. After the designated time, the plate was centrifuged for 2 min at 200 rcf at room temperature. The media was removed from each well and 200 μL of 5 μg/mL PI in 1x PBS solution. The plate was incubated for 30 minutes and again centrifuged. The fluorescence was read at the excitation wavelength of 535 nm and emission wavelength of 625 nm in a Biotek Synergy H1 monochromator.

### Preparation of extracellular vesicles

Media from the plated shRFP and shzDHHC21 control and necroptotic cells was collected and centrifuged at 300 rcf followed by 800 rcf for 30 mins at 4°C to remove cell debris. The supernatant was then filtered through a 0.2 μm filter followed by centrifugation at 100,000 rcf for 2h. The media was decanted and the pellet of extracellular vesicles was lysed followed by normalization based on protein content. The samples were then prepared for Western blotting.

### Generation of MLKL mutants

The point mutations in the pLys1-MLKL-Myc plasmid were conducted using the QuikChange II XL Site-Directed Mutagenesis Kit (Agilent Technologies Cat# 200521). XL-10 Gold ultracompetent cells were used for transformation and mutated plasmids were extracted using E.Z.N.A^®^ Plasmid DNA Mini Kit. The mutations were confirmed by Sanger sequencing using Genewiz (Azenta Life Sciences). The primers for introducing point mutations were designed on Benchling and obtained from Integrated DNA Technologies. Sequences of primers used to introduce point mutations are provided in Table S2.

### Acyl-Peg Exchange (APE)

APE procedure was modified based on previous protocols and has been described previously.^19^

### Digital droplet PCR (ddPCR)

Total RNA from shzDHHC5, shzDHHC20, shzDHHC21 and shRFP cells was extracted using the E.Z.N.A Total RNA kit. RNA was converted into cDNA using a Biorad Droplet Digital PCR QX200 system with a Biorad iScript cDNA synthesis kit. cDNA was diluted 1:20 for further use with molecular grade water. Reaction mixtures for ddPCR including the primers and cDNA were prepared with ddPCR Supermix for probes according to Biorad protocols and distributed in 96-well plates. A Biorad automated droplet generator was used to create water-oil emulsion droplets in a 96-well plate and subjected to PCR. The cycle was as follows: 95 ⁰C for 10 min, followed by 40 cycles of (1) 94 ⁰C for 30 s, (2) 56 ⁰C for 60 s, with a final 10-min inactivation step at 98 ⁰C. The plate was read using a QX200 droplet reader, which measured and quantified the fluorescence from the FAM fluorophore of the amplified target gene and the HEX fluorophore of the reference gene (HPRT1). Biorad Quantasoft software was used for data analysis and visualization of the data. The changes in gene expression and *p* values were calculated relative to HPRT1. The sequences for primers and probes are provided in Table S3.

### Western Blotting

Samples were loaded and separated with sodium dodecyl sulfate polyacrylamide gel (8% or 10%) electrophoresis at 150 V. Methanol activated polyvinyl difluoride (PVDF) membranes were used. The separated proteins were transferred onto the PVDF membranes at 50 V for 2 hours. Blocking conditions were incubation with 10% non-fat dry milk in tris-buffered saline (TBS)-Tween [10 mM Tris-base, 100 mM NaCl, 0.1% Tween 20 (pH 7.5)] at room temperature for 1 hour. Membranes were washed four times at 7 minute intervals in TBS-Tween. The corresponding membranes were incubated overnight at 4⁰C with primary antibodies (1:500/1:1000 for Myc-tag, 1:2000 dilution for MLKL, 1:500 for pMLKL, 1:10000 for flotillin 1, 1:10000 for calnexin, 1:5000 for α-tubulin). After incubation, the membranes were then washed four times with TBS-Tween for 10 minute each time. Dilutions for anti-rabbit and anti-mouse secondary antibodies were 1:2000 in 5% non-fat dry milk in TBS-Tween, followed by incubation at room temperature for one hour. The membranes were washed as earlier followed by developing with Super Signal West Pico kit.

### Quantification of western blots

Fiji software was used for the quantification of Western blots. Samples for all Western blot quantifications were loaded in triplicate. A frame for measurement was developed by using the rectangle tool of Fiji to cover the largest band of the protein of interest. The intensity of the other protein bands was similarly obtained. The background was also measured with the same frame to obtain background intensity measurements. The measured intensities from the protein of interest and background were then inverted by deducting measurements from 255. 255 is the pixel value assigned to white background. The inverted intensities of the protein bands of interest were corrected with the inverted intensities of the background. For APE blots, the relative intensities were obtained by dividing the corrected intensities of migrated bands by that of the non-migrated bands. For all other Western blots, the relative intensities were obtained by dividing the corrected intensity of MLKL or pMLKL to the corrected intensity of the loading control such as calnexin or tubulin. The calculations for every quantification have been described in the respective figure legends.

## QUANTIFICATION AND STATISTICAL ANALYSIS

All statistical analysis for the western blot quantifications, cell viability experiments and proteomics were performed using unpaired Student’s *t*-test. *p* values and numbers of replicates in all figures are indicated in figure legends, where *** *p* < 0. 001, ** *p* < 0.01, * *p* < 0.05, and ns is *p* > 0.05.

## Notes

### Competing Interest Statement

The authors have declared no competing interest.

### Summary of Updates

A revised version is uploaded.

