## Supplemental Information document for "Acylation of MLKL impacts its function in necroptosis"

**Figure S1. (A)** Overlay of the human and mouse MLKL with the cysteines in human MLKL highlighted in red with residue numbers in boxes above. The corresponding residues for mouse MLKL are marked in green. **(B)** Acyl PEG exchange with N-ethylmaleimide (NEM), hydroxylamine ($NH_2OH$) and methoxypolyethylene glycol maleimide (mPEG-Mal) was performed on control WT-MLKL and cysteine mutant cells. The conditions without $NH_2OH$ (represented by -) are controls for presence of acylation. The samples are blotted for myc. The control cells (not induced with necroptosis) show no distinguishable acylation/migration.

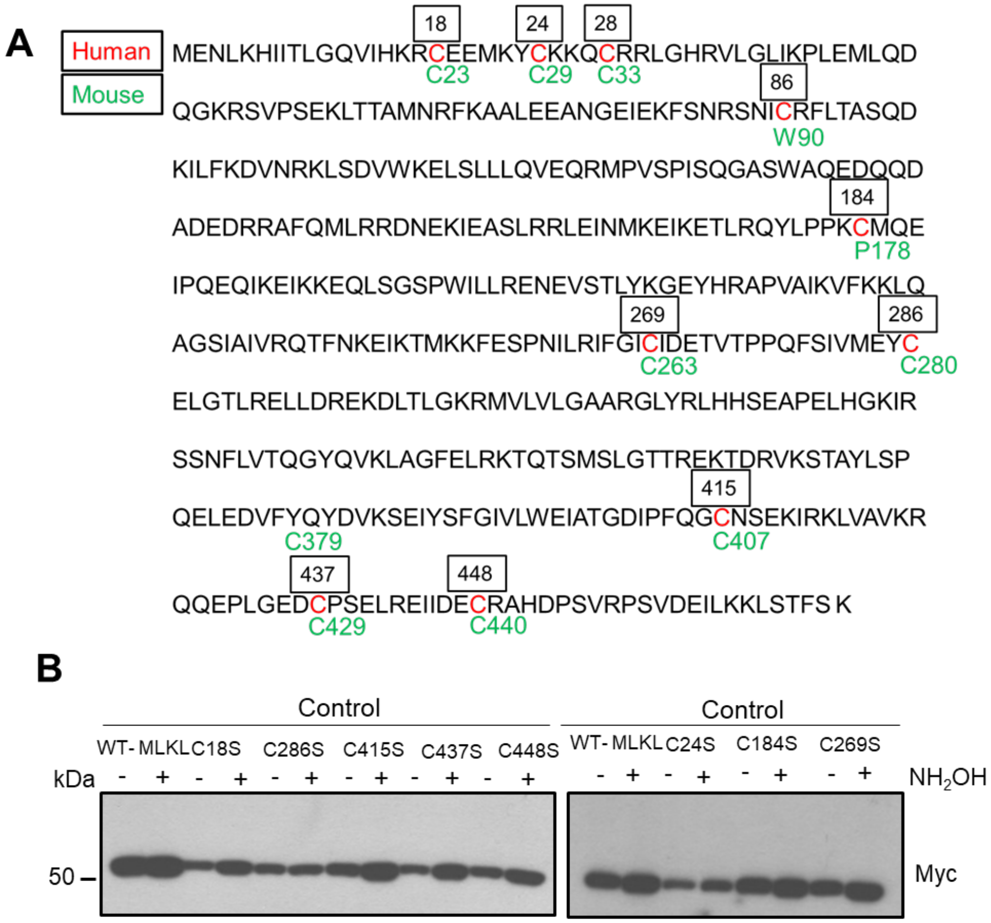

**Figure S2. (A)** Splay angles between fatty acid tails per lipid species in various simulation trajectories. Values reported correspond to the binding leaflet only. The splay angles were computed measuring the average angle between the vectors defined from the second carbon in each tail (C2) and the terminal carbon in the tail within 2.5nm of the protein. Splay angles per lipid species around the protein binding site in the binding and non-binding leaflets, respectively: **(B)** PIP, **(C)** $PIP_2$, **(D)** DOPC, **(E)** DOPE.

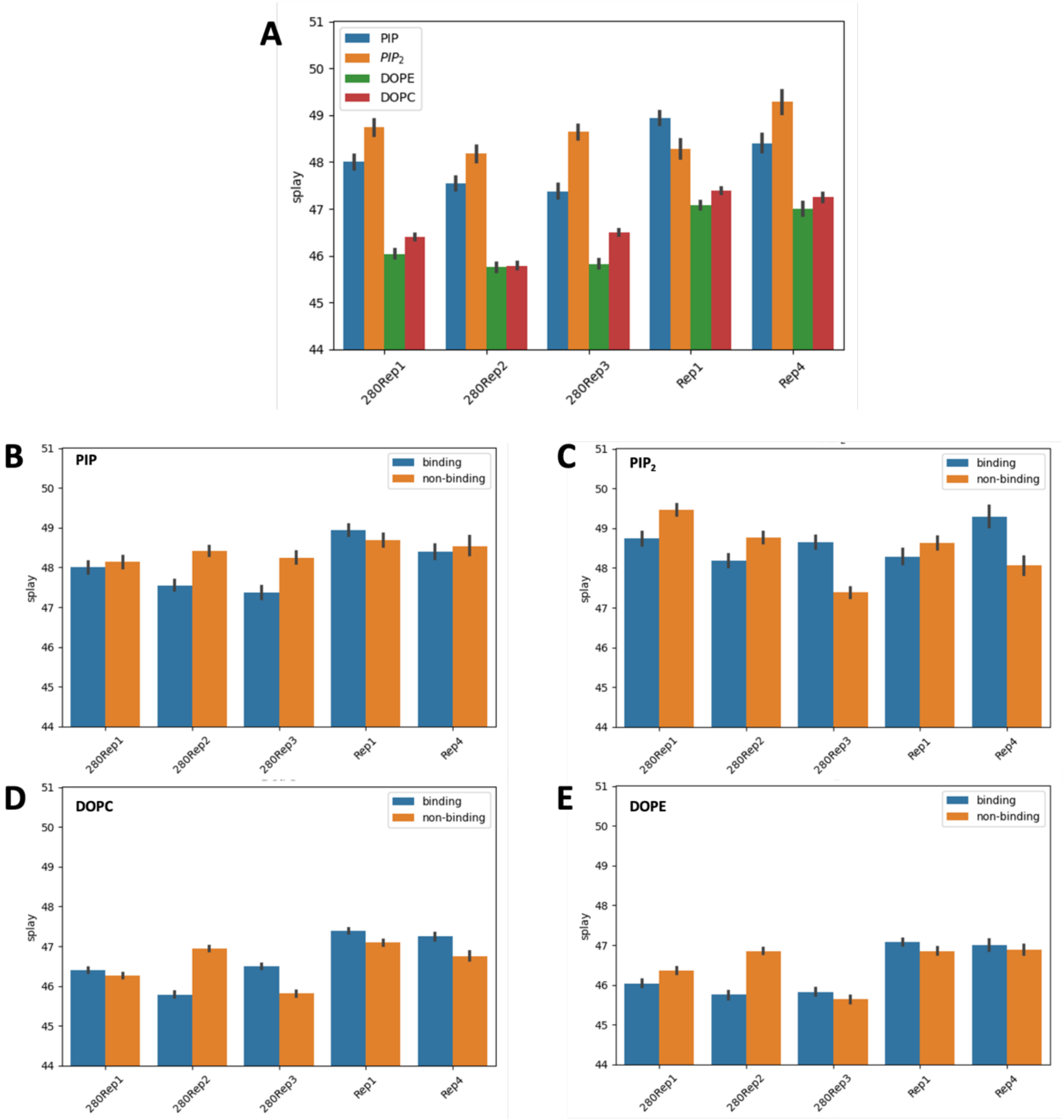

**Figure S3. (A)** Acyl PEG exchange with NEM, $NH_2OH$ and mPEG-Mal necroptotic shRFP and shzDHHC20 cells. The conditions without $NH_2OH$ (represented by -) are controls for presence of acylation. The samples are blotted for MLKL and pMLKL. **(B)** Quantification of acylated MLKL and pMLKL in shzDHHC20 compared to shRFP. The ratio is calculated by dividing migrated band intensities of pMLKL with non-migrated band intensities. No significant change in the extent of acylation between the two conditions was observed.

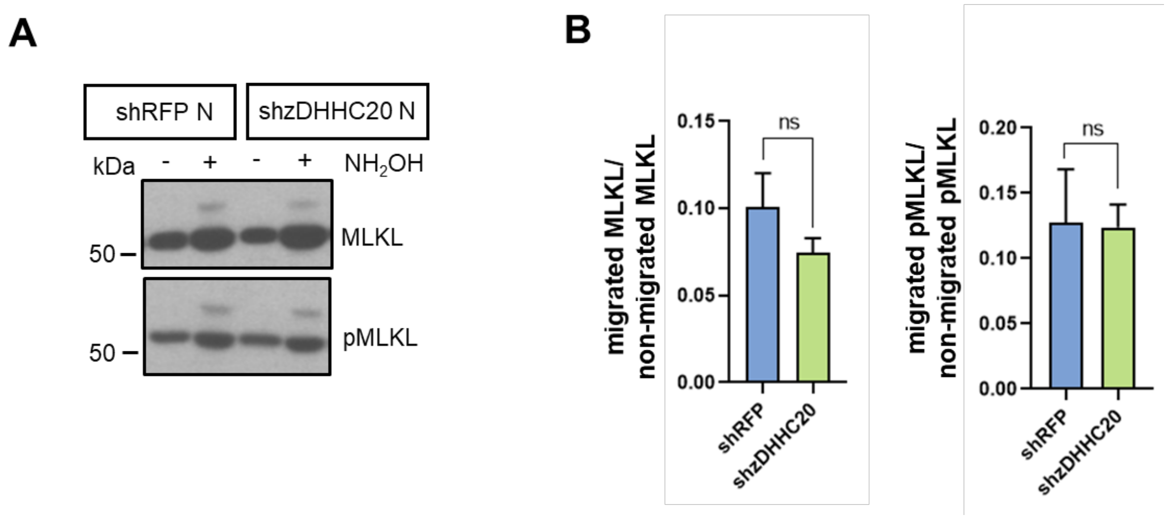

**Figure S4. (A)** Quantification showing shzDHHC21 has decreased pMLKL released in media in necroptosis. Media from control and necroptotic shRFP and shzDHHC21 cells was collected, filtered and subjected to ultracentrifugation to collect the extracellular vesicles and exosomes. The samples were normalized based on protein amounts and samples blotted for pMLKL. Bar plot represents relative protein levels calculated by normalizing the corrected intensities for each protein with the corresponding values for that protein for shRFP necroptotic condition. Data represent mean ± 1 SD; n=3. ** represents $p < 0.01$. **(B)** Quantification showing shzDHHC21 has decreased protein content released in media in necroptosis. Media from control and necroptotic shRFP and shzDHHC21 cells was collected, filtered and subjected to ultracentrifugation to collect the extracellular vesicles and exosomes. The samples were normalized based on protein amounts and samples blotted for FLOT1. Bar plot represents relative protein levels calculated by normalizing the corrected intensities for each protein with the corresponding values for that protein for shRFP necroptotic condition.

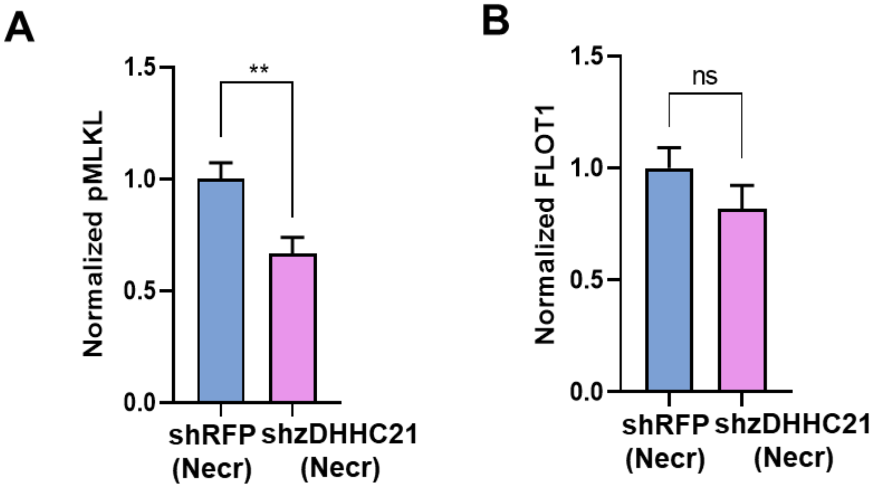

**Table S1.** Cysteine occupancies as percent of time each residue spent interacting with membrane lipids during the simulation. Cysteines of mouse MLKL presented with corresponding human MLKL residues shown in red. The trajectories for replicas 1, 2, 3 and 4 (Rep1, Rep2, Rep3 and Rep4) for the non-acylated MLKL have been previously published in Ramirez *et al*. (Ramirez et al.,2023)

| | | **C23**<br>**C18** | **C29**<br>**C24** | **C33**<br>**C28** | **C174**<br>**L180** | **C263**<br>**C269** | **C280**<br>**C286** | **C379**<br>**Y387** | **C407**<br>**C415** | **C429**<br>**C437** | **C440**<br>**C448** |
|---|---|---|---|---|---|---|---|---|---|---|---|
| **Non-acylated MLKL** | **Rep1** | 0.00 | 0.00 | 0.00 | 0.00 | 18.68 | 0.44 | 0.44 | **100.00** | 0.00 | 0.00 |
| | **Rep2** | **31.69** | 0.44 | 10.48 | 0.00 | 0.84 | 0.00 | **56.60** | 0.79 | 0.00 | 0.00 |
| | **Rep3** | **68.38** | 1.83 | 13.74 | 0.57 | 0.00 | 0.00 | 0.00 | 0.00 | 1.88 | 0.00 |
| | **Rep4** | 0.00 | 0.00 | 0.00 | 14.78 | 0.09 | **46.14** | 0.44 | **99.83** | 10.03 | 0.00 |
| **Acylated -MLKL** | **280r1**<br>**286** | 0.95 | 0.00 | 0.00 | 70.63 | 1.91 | 86.66 | 0.00 | 4.70 | 0.00 | 0.00 |
| | **280r2**<br>**286** | 0.00 | 0.00 | 0.00 | 1.72 | 5.34 | 60.17 | 0.00 | 83.97 | 0.10 | 0.00 |
| | **280r3**<br>**286** | 0.00 | 0.00 | 0.00 | 23.41 | 1.15 | 56.33 | 0.00 | 91.94 | 0.57 | 0.00 |

**Table S2.** Sequences of primers used to introduce point mutations:

| **Cysteine target** | **Primer sequence** |
|---|---|
| C18S | Forward: GTCATCCACAAACGGAGTGAAGAGATGAAATAC<br>Reverse: GTATTTCATCTCTTCACTCCGTTTGTGGATGAC |
| C24S | Forward: GAAGAGATGAAATACAGCAAGAAACAGTGCCGG<br>Reverse: CCGGCACTGTTTCTTGCTGTATTTCATCTCTTC |
| C184S | Forward: CAGTATTTACCACCAAAAAGCATGCAGGAGATCCCG<br>Reverse: CGGGATCTCCTGCATGCTTTTTGGTGGTAAATACTG |
| C269S | Forward: CTGCGTATATTTGGGATTAGCATTGATGAAACAGTGACTCCG<br>Reverse: CGGAGTCACTGTTTCATCAATGCTAATCCCAAATATACGCAG |
| C286S | Forward: ATTGTCATGGAGTACAGTGAACTCGGGACCCTG<br>Reverse: CAGGGTCCCGAGTTCACTGTACTCCATGACAAT |
| C415S | Forward: ATCCCGTTTCAAGGCAGTAATTCTGAGAAGATC<br>Reverse: GATCTTCTCAGAATTACTGCCTTGAAACGGGAT |
| C437S | Forward: CCACTGGGTGAAGACAGCCCTTCAGAGCTGCGG<br>Reverse: CCGCAGCTCTGAAGGGCTGTCTTCACCCAGTGG |
| C448S | Forward: GAGATCATTGATGAGAGCCGGGCCCATGATCCC<br>Reverse: GGGATCATGGGCCCGGCTCTCATCAATGATCTC |

**Table S3.** Sequences of primers used for droplet digital PCR:

| Target | Primer and probe sequence |
|---|---|
| zDHHC5 | Forward: CAGGGAAAGGAGGAAAAGGAA<br>Reverse: GTCTGTACAACTGTGTGGAG<br>Probe: 56-FAM/AACTGTATT/ZEN/GGTCGCCGGAACTACC/3IABkFQ |
| zDHHC20 | Forward: CCATTTCCATCAGGTCCGTAT<br>Reverse: GTTCTTCATCAGCGTCCTCTC<br>Probe: 56-FAM/TTCCAACTA/ZEN/GCCAGCAGTGGTAGC/3IABkFQ |
| zDHHC21 | Forward: CACTTGTTACATAATTCCCAGAACT<br>Reverse: GGCCTCCATAACTGATCCAG<br>Probe: 56-FAM/CCTGAGAAC/ZEN/CCCAAGATCCCACAT/3IABkFQ |
| HPRT1 | Forward: GTATTCATTATAGTCAAGGGCATATCC<br>Reverse: AGATGGTCAAGGTCGCAAG<br>Probe: 5HEX/TGGTGAAAA/ZEN/GGACCCCACGAAGT/3IABkFQ/ |
